# Endogenous autophagosome reporters reveal redundant and non-redundant roles of LGG-1 and LGG-2 in *C. elegans* neuronal autophagy

**DOI:** 10.64898/2026.07.15.738464

**Authors:** Heather Tsong, Mya Rodriguez, Andrea K. H. Stavoe

**Author notes:** These authors contributed equally (Heather Tsong, Mya Rodriguez).

## Abstract

Autophagy is a conserved cellular recycling pathway essential for neuronal development and homeostasis. Neurons are highly polarized cells that rely on the continual turnover of cytoplasmic content through processes like autophagy to maintain cellular function. Here, we employed novel endogenous autophagosome reporters to define the distinct expression patterns, spatial distribution, and compensatory potential of the two *C. elegans* Atg8 orthologs, LGG-1 and LGG-2, in the nervous system. We labelled LGG-1 and LGG-2 with split wrmNeonGreen to endogenously label autophagosomes in individual neurons. We observed brighter and larger wrmNG_split_:LGG-1 autophagosomes relative to smaller and dimmer wrmNG_split_:LGG-2 autophagosomes pan-neuronally and in the individual AIY and NSM neurons. When we interrogated autophagosome trafficking in the AIY neurite, we revealed a similar trafficking mechanism for both LGG-1 and LGG-2 puncta. In AIY and pan-neuronally, we found that *lgg-2* was required for LGG-1 puncta formation. Strikingly, in *lgg-1* mutants, LGG-2 demonstrated a compensatory capacity in the AIY neuron via an upregulation of LGG-2 autophagosomes. Overall, our findings demonstrate the advantages of this novel endogenous reporter system for tracking neuronal autophagy and further elucidate redundant and non-redundant functions of LGG-1 and LGG-2.

## INTRODUCTION

Macroautophagy, hereafter autophagy, is a highly conserved cellular recycling pathway. Autophagy supports cellular homeostasis by engulfing intracellular components, such as protein aggregates and organelles, within a double-membrane vesicle called the autophagosome. The autophagosome delivers these sequestered materials to the lysosome for degradation and recycling. During autophagosome biogenesis, the sequential action of multiple protein complexes leads to the formation of a cup-shaped double membrane called the phagophore. The phagophore membrane is then extended and eventually closes to form the autophagosome (Klionsky and Emr 2000; Glick et al. 2010; Melia et al. 2020). Atg8, a ubiquitin-like protein that associates with the autophagosomal membrane, is essential for autophagosome formation. As the phagophore membrane extends, Atg8 is processed and conjugated to phosphatidylethanolamine (PE) on the inner and outer leaflet of the phagophore through a ubiquitination-like cascade. Lipidation of Atg8 is necessary for autophagosome biogenesis to proceed; however, the exact functions of PE-conjugated Atg8 are unclear. Lipidated Atg8 appears to be involved in many steps of autophagy, including phagophore membrane extension, membrane tethering, cargo recognition, autophagosome transport, and lysosomal fusion (Shpilka et al. 2011; Lystad and Simonsen 2019). Atg8 and its orthologs are commonly used as protein markers of autophagosomes because Atg8 associates with the phagophore early in autophagosome biogenesis and remains attached to the membrane until autophagosomal contents are degraded by the lysosome (Klionsky et al. 2021).

Atg8 is highly conserved from yeast to nematodes to mammals. While yeast have one Atg8 gene, mammals have five or six (mAtg8s). The mAtg8 family can be divided into subfamilies based on sequence similarity: the LC3 family, which includes LC3A, LC3B, and LC3C, and the GABARAP family, which includes GABARAP, GABARAPL1, and GABARAPL2/GATE-16 (Shpilka et al. 2011; Varga et al. 2022). *Caenorhabditis elegans* have two Atg8 orthologs, with one representative in each mAtg8 subfamily: “LC3, GABARAP and GATE-16 family” *lgg-1* and *lgg-2*. LGG-1 is orthologous to the GABARAP subfamily and LGG-2 is more similar to the LC3 subfamily (Fig. 1A) (Jenzer et al. 2014; Manil-Ségalen et al. 2014).

**Figure 1.**
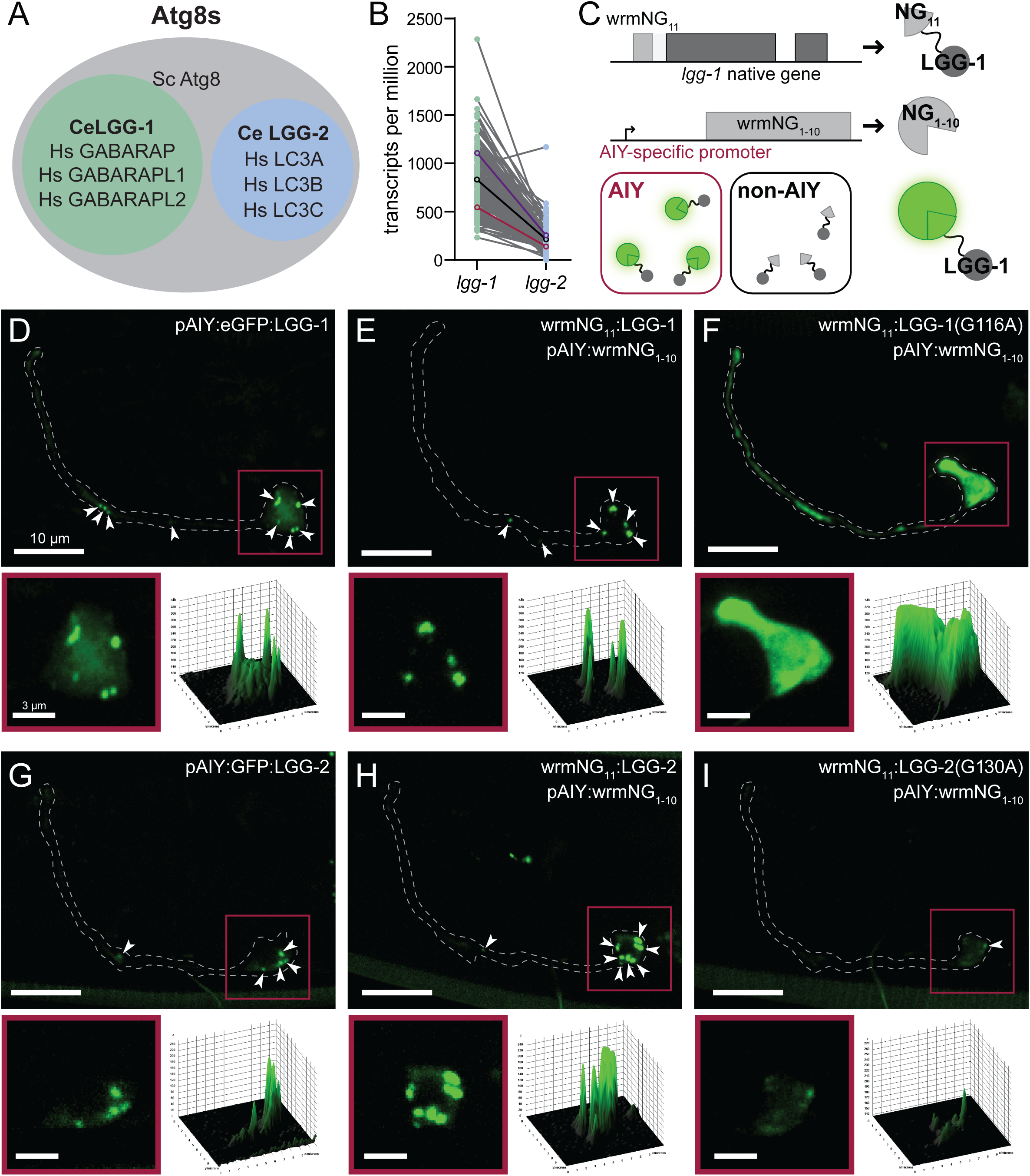
Endogenous labeling of LGG-1 and LGG-2 in AIY. (A) Atg8 subfamilies. Hs: Homo sapiens; Ce: Caenorhabditis elegans; Sc: Saccharomyces cerevisiae. (B) Expression of *lgg-1* (green) and *lgg-2* (blue) in *C. elegans* neurons. The expression levels for a given neuron are connected by a gray line. Black dots and line represent nervous system mean. Red identifies AIY and purple identifies NSM in data set. Data from (Taylor et al. 2021). (C) Schematic of split-wrmNG endogenous labeling strategy for LGG-1 in AIY. (D-I) Representative maximal projection micrographs of z-stacks of whole AIY (top) and AIY soma (below: inset 3D surface plot) of: ectopically expressed eGFP:LGG-1 (D), endogenously labeled wrmNG_split_:LGG-1 (E), endogenously expressed lipidation mutant LGG-1(G116A) (F), ectopically expressed GFP:LGG-2 (G), endogenously labeled wrmNG_split_:LGG-2 (H), and endogenously expressed lipidation mutant LGG-2(G130A) (I). Inset is indicated by red box in top micrographs. Arrowheads indicate LGG puncta. AIY boundary is indicated by dashed white line. Scale bars, 10 μm (inset scale bars, 3 μm). Micrographs represented in D-E were acquired at 90 ms exposure, while G-I were acquired at 400 ms exposure.

Like in yeast, mammalian and worm Atg8s are processed and conjugated to PE at the phagophore membrane (Zhang et al. 2015; Klionsky et al. 2021). However, the presence of multiple Atg8 genes in multicellular organisms begs the question: are there functional differences between them? Studies in mammalian cell culture suggest that, while both LC3 and GABARAP family proteins are necessary for autophagy, they act at distinct steps in autophagosome biogenesis (Weidberg et al. 2010; Nguyen et al. 2016; Bui et al. 2024). They may also be differentially utilized in specific forms of selective autophagy (Johansen and Lamark 2020). Although evidence suggests distinct functions for the mAtg8s, LC3B is the most commonly used marker for autophagosomes in mammals (Klionsky et al. 2021). Interestingly, LGG-1, which is more closely related to mammalian GABARAP, is most commonly used as an autophagosome marker in *C. elegans* in lieu of LGG-2, which is more closely related to LC3 (Manil-Ségalen et al. 2014; Zhang et al. 2015). In *C. elegans*, lgg-1 and lgg-2 appear to have both redundant and non-redundant functions, depending on the context (Manil-Ségalen et al. 2014; Wu et al. 2015; Leboutet et al. 2023). Previous studies have sought to dissect these differences; however, few have probed endogenous proteins in live animals.

Much of what we know about the molecular mechanisms of autophagosome biogenesis comes from studies performed in yeast and immortalized mammalian cell lines, particularly in response to acute stressors such as starvation (Glick et al. 2010; Hale et al. 2013; Abada and Elazar 2014). However, previous work has demonstrated that neurons regulate autophagy differently than non-polarized cells (Tsvetkov et al. 2010; Maday and Holzbaur 2016). Constitutive, nonselective autophagy functions at a high, constant level in neurons *in vitro* and *in vivo* (Boland et al. 2008). Neuronal autophagosome biogenesis is also spatially regulated, with autophagosome formation occurring predominantly in the distal axon. Cytoplasmic dynein then transports autophagosomes back to the cell body; during retrograde transport, autophagosomes mature and fuse with lysosomes (Maday et al. 2012; Maday and Holzbaur 2014; Soukup et al. 2016; Stavoe et al. 2016). Importantly, these studies interrogating the spatial and temporal kinetics and mechanisms of neuronal autophagy were conducted either in fixed samples or with ectopically expressed tagged proteins.

In the autophagy field broadly, autophagosomes have historically been visualized by exogenously introduced tagged Atg8s or by immunostaining of fixed cells or tissues (Klionsky et al. 2021). A ubiquitously used transgenic mouse ectopically expresses GFP:LC3B (Mizushima et al. 2004). In *C. elegans* specifically, autophagosomes are more commonly visualized with exogenous expression of tagged LGG-1 (Zhang et al. 2015; Palmisano and Meléndez 2016; Chang et al. 2017; Klionsky et al. 2021). While these methods have yielded important results, there are limitations. Exogenous transgenes are highly overexpressed and are not governed by the same regulatory mechanisms that the endogenous gene is subjected to. In *C. elegans*, transgenes are typically delivered through gonadal microinjections and are inherited as extrachromosomal arrays. How these extrachromosomal arrays assemble and how many copies are present depend on a multitude of factors, many of which cannot be experimentally controlled. As a result, there is variability among the reporter strains depending on how they were generated. Integrated transgenes offer greater stability but remain overexpressed (Mello and Fire 1995; El Mouridi et al. 2022). Endogenous proteins can be probed by immunostaining, but this must be performed on fixed tissues, precluding any dynamic experiments, and requires reliable antibodies.

Endogenous tagging of target proteins in live worms is now achievable with genome editing via CRISPR/Cas9 (Cho et al. 2013; Arribere et al. 2014; Zhao et al. 2014; Paix et al. 2015; Prior et al. 2017; Dokshin et al. 2018). While one group has endogenously tagged LGG-1, their strategy relied on the use of a full-length fluorophore inserted into the endogenous *lgg-1* locus (Bördén et al. 2025), making cell-or tissue-specific visualization of autophagosomes challenging, particularly in the cramped quarters of the nervous system. Here, we have generated a novel, endogenous autophagosome reporter using CRISPR/Cas9 to visualize autophagosome biogenesis in single tissues or cells without overexpression. We used this system to interrogate the differences between LGG-1 and LGG-2 in neuronal autophagy in live animals.

## RESULTS

### Visualizing autophagosomes in a single neuron with endogenous tagging of LGG-1 and LGG-2

We set out to interrogate the autophagy pathway at the level of individual neurons within the context of intact *Caenorhabditis elegans* so that we could observe autophagosomes marked with endogenously labelled proteins. We designed a system to flexibly drive visualization of this autophagosome reporter in individual neurons with neuron-specific promoters (Brenner 1974; Flibotte et al. 2010). We chose to use the split mNeonGreen (NG) system. mNG is over three times brighter than GFP in *C. elegans*, making it ideal for endogenous labeling (Hostettler et al. 2017). We translated the optimized split mNG (Feng et al. 2017; Zhou et al. 2020) into *C. elegans* (wrmNG) by codon-optimizing each part of the split mNG for worms. To label autophagosomes, we started by tagging *lgg-1* because *lgg-1* is more highly expressed than *lgg-2* throughout the nervous system (Fig. 1B) (Taylor et al. 2021). Modifying the Native and Tissue-Specific Fluorescence strategy (He et al. 2019), we inserted the 11^th^ beta-strand of wrmNG (wrmNG_11_) on the N-terminal side of LGG-1 using CRISPR/Cas9. We drove expression of the rest of the fluorophore (wrmNG_1-10_) by pan-neuronal or neuron-specific promoters; this construct also included a transcriptionally coupled mScarlet to mark target cells independent of endogenous LGG-1 levels. Thus, full-length, fluorescent wrmNG was only reconstituted in tissues or cells that expressed both LGG-1 and the exogenous array (Fig. 1C). As wrmNG_1-10_ is overexpressed, the limiting factor to signal brightness is the expression level of *lgg-1*. For ease of understanding, we will refer to the endogenously tagged LGG-1 construct as wrmNG_split_:LGG-1.

Autophagosomes were previously studied in interneuron AIY (Amphid Interneuron Y) using ectopically expressed GFP:LGG-1 (Stavoe et al. 2016; Hill et al. 2019). Using previously generated worms that ectopically expressed eGFP:LGG-1 in AIY, we observed autophagosomes in the AIY cell body and neurite, consistent with previous studies (Fig. 1D). We next drove wrmNG_1-10_ expression in AIY in worms harboring endogenously tagged *wrmNG_11_:lgg-1*. We observed wrmNG fluorescent signal only in AIY (Fig. 1E). The endogenously tagged wrmNG_split_:LGG-1 revealed discrete, punctate structures in the AIY neurite and soma (Fig. 1E). In contrast to ectopically expressed eGFP:LGG-1, endogenously tagged wrmNG_split_:LGG-1 had little to no cytoplasmic background signal in AIY, allowing for easy identification of puncta (Fig. 1D-E).

Next, to verify that the endogenous tag does not interfere with LGG-1 localization, we leveraged the *lgg-1(G116A)* mutation. In this mutant, glycine 116 is mutated to alanine, preventing LGG-1 cleavage by ATG-4, which is necessary for LGG-1 lipidation to PE on the phagophore (Mizushima et al. 2010; Zhang et al. 2015; Chen et al. 2017). Endogenous wrmNG_split_:LGG-1(G116A) signal was cytoplasmic instead of localized to discrete puncta (Fig. 1F, Fig. S1A), validating that the endogenous tag does not interfere with LGG-1 localization (Fig. 1E). Interestingly, wrmNG_split_:LGG-1(G116A) signal was extremely bright (Fig. S1B), suggesting a build-up of mutant LGG-1 that is no longer turned over by canonical autophagy. Together, our data suggest that we have successfully tagged endogenous LGG-1 without disrupting its localization to autophagosomal structures, validating our split-wrmNG endogenous LGG-1 system.

Since worms have two Atg8 orthologs, we next asked if we could endogenously tag LGG-2 in a similar manner. We employed the same strategy for endogenous tagging of LGG-2. Consistent with our LGG-1 data and previous studies, exogenously tagged LGG-2 forms puncta in the AIY cell body and neurite (Fig. 1G). Endogenously tagged wrmNG_split_:LGG-2 also localizes to punctate structures in the AIY neurite and soma (Fig. 1H). Compared to extrachromosomal eGFP:LGG-2, endogenous wrmNG_split_:LGG-2 is brighter, although there is still some cytoplasmic signal present in both tagging strategies (Fig. 1G-H). As before, we validated the endogenous tag using an LGG-2 lipidation mutant. Cleavage of LGG-2 by ATG-4 occurs at glycine 130 and mutating it to alanine prevents LGG-2 cleavage and lipidation to PE (Alberti et al. 2010; Manil-Ségalen et al. 2014; Chen et al. 2017). We mutated the endogenously tagged LGG-2 to LGG-2(G130A) using CRISPR genome editing. Similar to our results with wrmNG_split_:LGG-1(G116A), we observed fewer wrmNG_split_:LGG-2(G130A) puncta and increased cytoplasmic signal compared to wild-type eGFP:LGG-2 (Fig. 1I). In contrast to our results with wrmNG_split_:LGG-1(G116A), we observed a substantial reduction in wrmNG_split_:LGG-2(G130A) overall signal. These data validate our endogenously tagged LGG-2 marker. Taken together, our results indicate that we can visualize endogenously tagged LGG-1 and LGG-2 in a single neuron using the split-NeonGreen system. Furthermore, endogenous wrmNG_split_:LGG-1 and wrmNG_split_:LGG-2 display less background than overexpressed tags, making automated downstream analysis of LGG-1 and LGG-2 puncta more feasible.

### LGG-1 and LGG-2 puncta at the pan-neuronal and single neuron level

We next examined differences between endogenous LGG-1 and LGG-2 expression at the pan-neuronal level using a rab-3 promoter (Fig. 2A). In contrast to the extrachromosomal array expression of wrmNG_1-10_ in AIY (Fig. 1), we used the SKI LODGE system (Silva-García et al. 2019) to generate a single-copy cassette driving pan-neuronal expression of wrmNG_1-10_ and cytoplasmic wrmScarlet (wrmScar). We visualized wrmNG_split_:LGG-1 and wrmNG_split_:LGG-2 independently in all *C. elegans* neurons. To quantify autophagosomes in the worm nervous system, we focused on neuronal cell bodies posterior to the nerve ring (Fig. 2B, 2D). We developed an automated pipeline using ImageJ 3D Objects Counter to identify puncta within the specified volume (Fig. 2C) and quantify the number, mean fluorescence intensity, and volume of wrmNG_split_:LGG-1 and wrmNG_split_:LGG-2 puncta. We observed more wrmNG_split_:LGG-2 than wrmNG_split_:LGG-1 puncta (Fig. 2E). This was unexpected, as previous single-cell transcriptomic analyses in *C. elegans* has shown that *lgg-*1 is more highly expressed than *lgg-2* across neurons (Taylor et al. 2021). However, further analysis showed that wrmNG_split_:LGG-1 puncta were brighter and larger than wrmNG_split_:LGG-2 puncta (Fig. 2F-G), consistent with higher levels of *lgg-1* expression. Lower expression levels of *lgg-2* likely allowed the automated pipeline to more easily distinguish discrete wrmNG_split_:LGG-2 puncta. In contrast, higher expression of *lgg-1* did not allow the pipeline to distinguish between the brighter wrmNG_split_:LGG-1 puncta, resulting in seemingly fewer wrmNG_split_:LGG-1 puncta. Taken together, these results show that there is more endogenous LGG-1 than LGG-2 expression at the pan-neuronal level.

**Figure 2.**
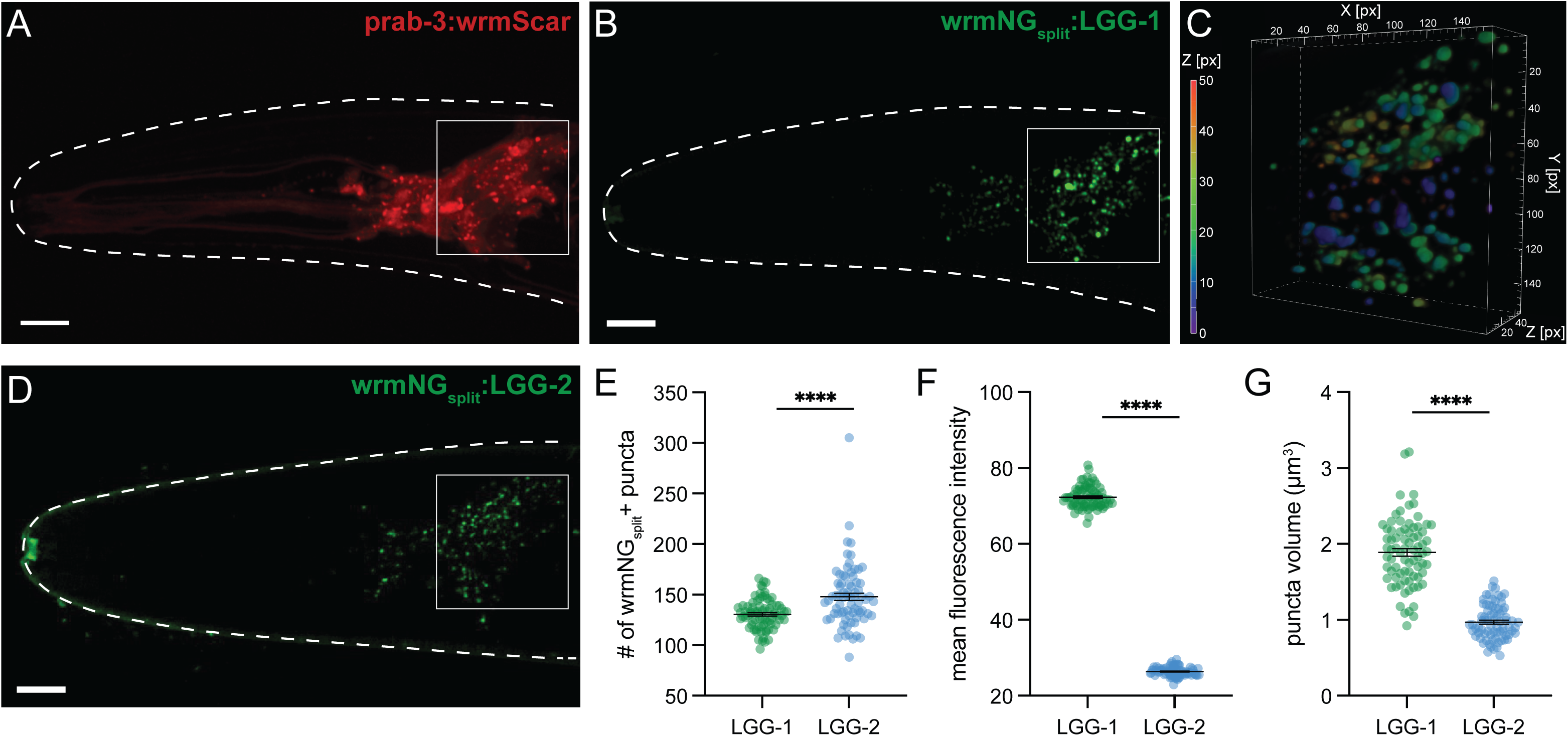
Endogenously labeled LGG-1 is brighter than LGG-2 in the nerve ring. (A) Representative maximal projection micrograph of a z-stack of transcriptionally coupled wrmScarlet visualizing where wrmNG_1-10_ is expressed in the head of the worm. (B) Representative maximal projection micrograph of endogenously labeled wrmNG_split_:LGG-1 in the head. (C) Representative volumetric image of wrmNG_split_:LGG-1 puncta within region of interest indicated by the white boxes in A and B. (D) Representative maximal projection micrograph of endogenously labeled wrmNG_split_:LGG-2 in the head of the worm. Anterior is to the left. Scale bars, 10 μm. (E-G) Quantification of the number of puncta (E), the mean fluorescence intensity of puncta (F) and the puncta volume (G) of endogenously labeled LGG-1 and LGG-2 in the nerve ring. Mean ± SEM. **** p<0.0001 by Mann Whitney U test. Location of volume analyzed is depicted by white box in A, B, and D.

Next, we quantified and compared wrmNG_split_:LGG-1 and wrmNG_split_:LGG-2 puncta at the single-neuron level in AIY (Fig. 3A-C). We further stratified our analysis between the soma and the neurite. In the AIY soma, wrmNG_split_:LGG-1 puncta were more abundant (Fig. 3D) and brighter (Fig. 3E) than wrmNG_split_:LGG-2 puncta; however, wrmNG_split_:LGG-1 and wrmNG_split_:LGG-2 puncta were of similar size (Fig. 3F). In the AIY neurite, wrmNG_split_:LGG-1 puncta were more abundant (Fig. 3G), brighter (Fig. 3H), and larger (Fig. 3I) than wrmNG_split_:LGG-2 puncta. These results are broadly consistent with our pan-neuronal results but suggest that we can more readily distinguish between wrmNG_split_:LGG-1 puncta in a single neuron compared to the entire nerve ring.

**Figure 3.**
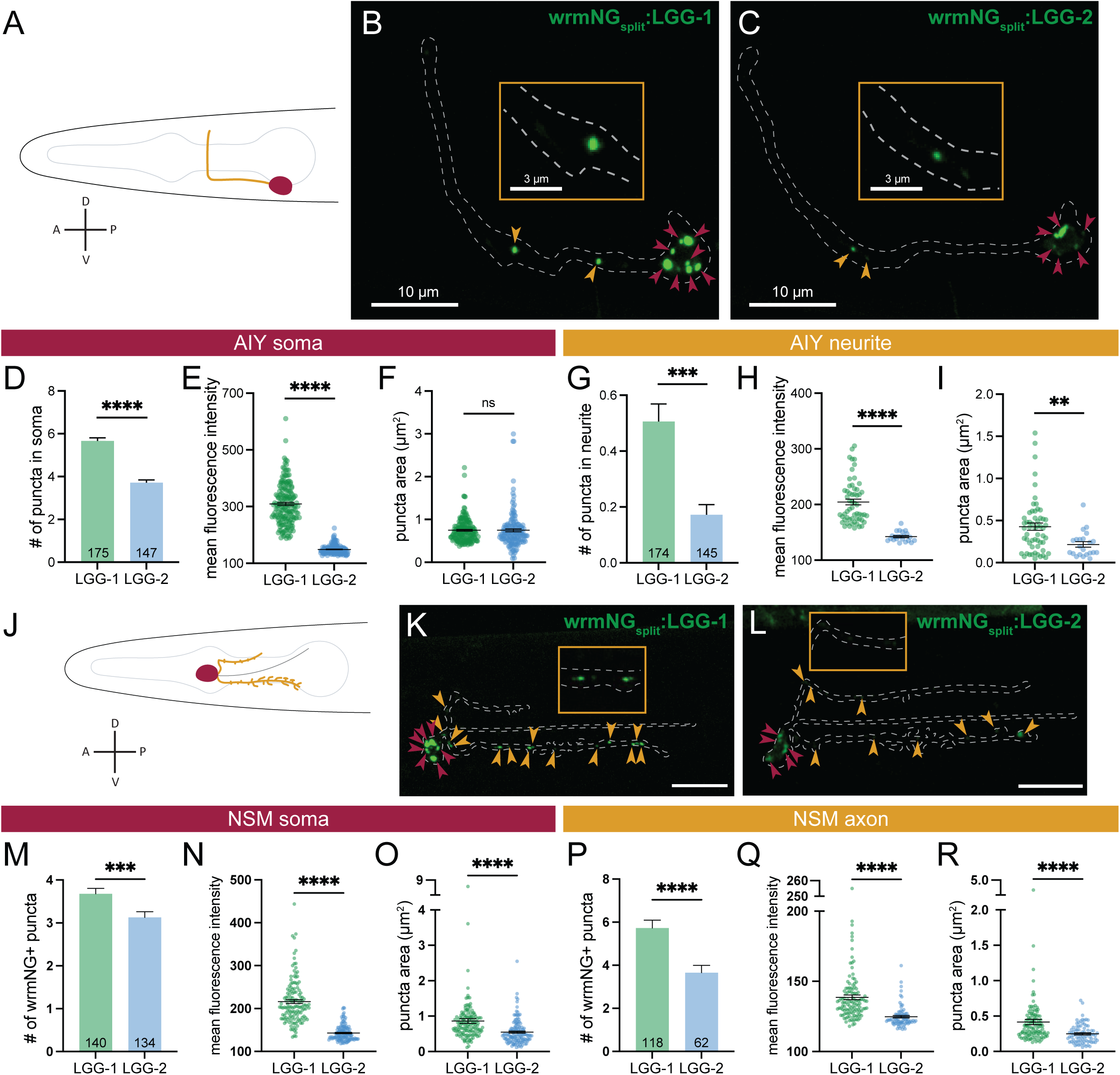
LGG-1 puncta are brighter, larger and more numerous than LGG-2 puncta in AIY and NSM. (A) Schematic of AIY in the worm head. (B-C) Representative maximal projection micrographs of z-stacks of endogenously labeled wrmNG_split_:LGG-1 (B) and wrmNG_split_:LGG-2 (C) in AIY. (D-I) Quantifications in the AIY soma (D-F) and AIY neurite (G-I) of puncta number (D, G), mean fluorescence intensity (E, H) and puncta area (F, I) for LGG-1 and LGG-2 puncta. Mean + or ± SEM. (J) Schematic of NSM in the worm head. (K-L) Representative maximal projection micrographs of z-stacks of endogenously labeled wrmNG_split_:LGG-1 (K) and wrmNG_split_:LGG-2 (L) in NSM. (M-R) Quantifications in the NSM soma (M-O) and NSM axons (P-R) of puncta number (M, P), mean fluorescence intensity (N, Q) and puncta area (O, R) for LGG-1 and LGG-2 puncta. In A and J, A is anterior, D is dorsal, P is posterior and V is ventral. In B, C, K and L, red arrowheads indicate puncta in soma, gold arrowheads indicate puncta in neurites. Insets magnify presynaptic regions in AIY and NSM. Neuron boundary is indicated by dashed white line. In D, G, M and P, n values are indicated at the bottom of each bar in the graph; in other graphs, all values are plotted. **** p < 0.0001; *** p < 0.0005; ** p < 0.005; ns p> 0.05 by Welch’s t test (D) or Mann-Whitney test (E-I, M-R).

We next investigated whether we could visualize autophagosomes with these endogenous markers in a different neuron, NSM (Fig. 3J-L). NSM (<u>N</u>euro<u>S</u>ecretory <u>M</u>otor) has a cell body with a bifurcated axon and a dendrite (Albertson and Thompson 1976; Axäng et al. 2008). The subventral branch of the axon is the longest and exhibits complex axonal arbors while the dorsal axon is short with little arborization (Nelson and Colón-Ramos 2013). Similar to AIY, we observed higher endogenous expression of wrmNG_split_:LGG-1 versus wrmNG_split_:LGG-2 in NSM. In the NSM soma, we observed wrmNG_split_:LGG-1 puncta that were more abundant (Fig. 3M), brighter (Fig. 3N), and larger (Fig. 3O) compared to wrmNG_split_:LGG-2 puncta. Congruently, in the NSM axon, wrmNG_split_:LGG-1 puncta were also more abundant (Fig. 3P), larger (Fig. 3Q), and brighter (Fig. 3R) than wrmNG_split_:LGG-2 puncta. We did not observe any wrmNG_split_:LGG-1 or wrmNG_split_:LGG-2 puncta in the NSM dendrite. Taken together, our data indicate that wrmNG_split_:LGG-1 puncta are more numerous, brighter, and larger than wrmNG_split_:LGG-2 puncta in neurons broadly and specifically in AIY and NSM.

### Analogous trafficking of LGG-1 and LGG-2-labelled autophagosomes in AIY

In neurons, after formation in the axon, autophagosomes are retrogradely trafficked along microtubules by cytoplasmic dynein to the cell body. As they approach the soma, autophagosomes fuse with endosomes and lysosomes for degradation and cargo recycling (Maday et al. 2012; Stavoe et al. 2016). UNC-16/JIP3 regulates this retrograde transport of autophagosomes by acting as an activating adaptor for dynein (Cason and Holzbaur 2023). Thus, mutations to *unc-16* prevent autophagosome retrograde trafficking. Given the observed differences between LGG-1 and LGG-2 puncta, we next asked whether LGG-1 and LGG-2 puncta are formed and trafficked similarly. Previous studies indicate that in AIY, autophagosomes form in the neurite at presynaptic sites (Stavoe et al. 2016; Hill et al. 2019). In *unc-16* mutant animals, ectopically labelled GFP:LGG-1 puncta aberrantly accumulate in the neurite, as the autophagosomes are not delivered to the soma and are not turned over through lysosomal fusion (Hill et al. 2019). Thus, mutations to *unc-16* prevent autophagosome retrograde trafficking.

We were able to replicate the previously described phenomenon using endogenously tagged wrmNG_split_:LGG-1. In *unc-16* mutants, we observed thirteen times more wrmNG_split_:LGG-1 puncta in the AIY neurite as compared to wild-type animals (Fig. 4A-C). In contrast, we observed no significant change to the number of wrmNG_split_:LGG-1 puncta in the soma (Fig. 4D), suggesting that autophagosome biogenesis occurring in the soma is unaffected in *unc-16* mutants. We then examined wrmNG_split_:LGG-2 puncta in *unc-16* mutant animals, similarly observing a significant accumulation of wrmNG_split_:LGG-2 puncta in AIY neurites (Fig. 4E-G). Once again, the number of wrmNG_split_:LGG-2 puncta in the soma did not change in *unc-16* mutant animals (Fig. 4H). While there was an accumulation of both wrmNG_split_:LGG-1 and wrmNG_split_:LGG-2 puncta in the AIY neurites of *unc-16* mutant animals, the relative characteristics of these puncta did not change. In *unc-16* mutants, wrmNG_split_:LGG-1 puncta were still more abundant (Fig. 4I), brighter (Fig. 4J), and larger (Fig. 4K) than wrmNG:LGG-2 puncta. These data are consistent with studies in primary mammalian neurons and suggest that both LGG-1 puncta and LGG-2 puncta are generated in the neurite and that their retrograde trafficking is regulated by UNC-16/JIP3.

**Figure 4.**
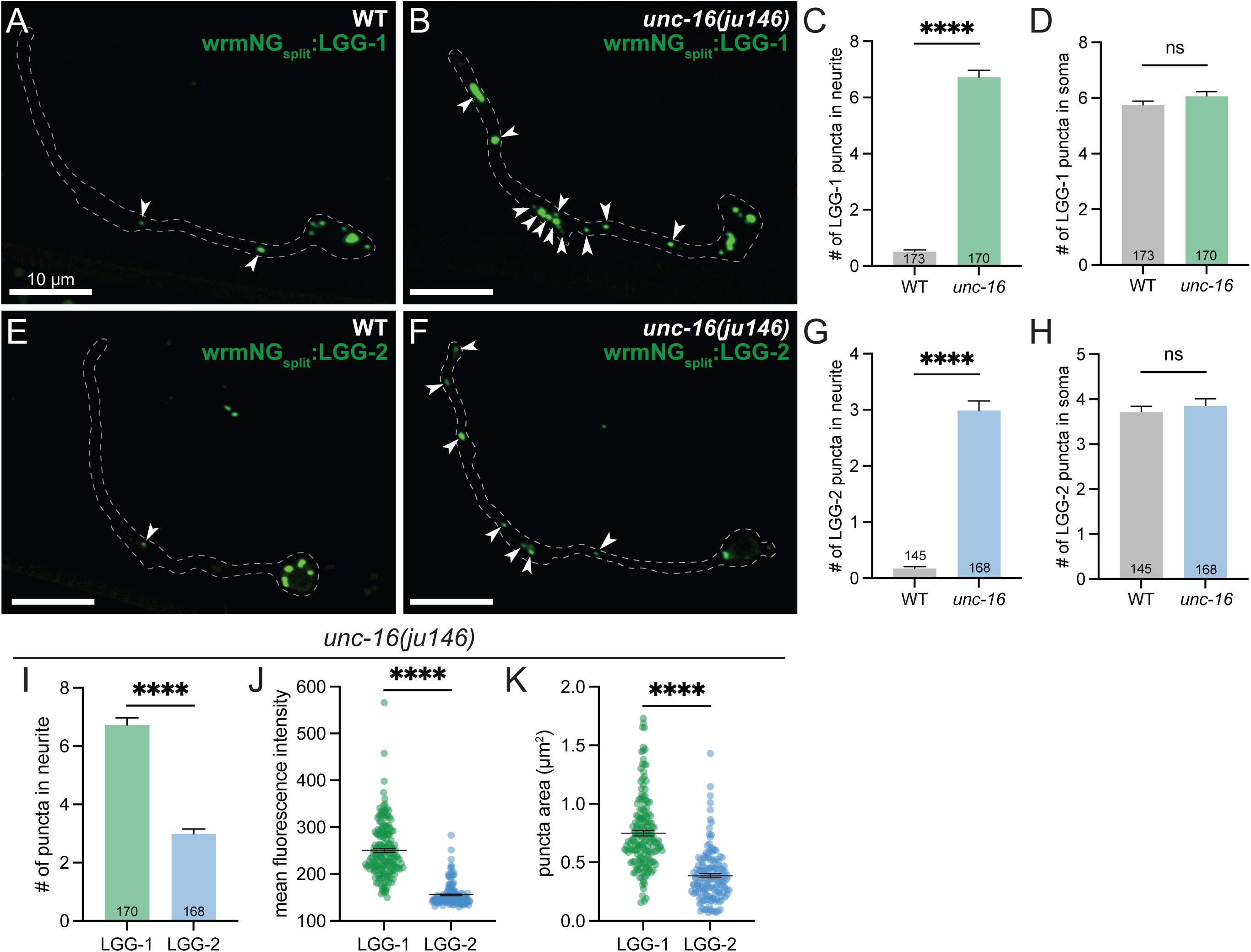
UNC-16/JIP3 is necessary to transport LGG-1 and LGG-2 puncta. (A-B) Representative maximal projection micrographs of z-stacks of endogenously labeled wrmNG_split_:LGG-1 in AIY in wild type (A) and *unc-16* (B) mutant worms. (C-D) Quantification of LGG-1 puncta number in AIY neurite (C) and soma (D). Mean + SEM. (E-F) Representative maximal projection micrographs of z-stacks of endogenously labeled wrmNG_split_:LGG-2 in AIY in wild type (E) and *unc-16* (F) mutant worms. (G-H) Quantification of LGG-2 puncta number in AIY neurite (G) and soma (H). Mean + SEM. (I-K) Quantifications of puncta number, mean fluorescence intensity and puncta area for LGG-1 and LGG-2 puncta in *unc-16* mutant worms. Mean + or ± SEM. In C, D, G, H and I, n values are indicated at the bottom of each bar in the graph; in other graphs, all values are plotted. **** p<0.0001; ns p > 0.05 by Welch’s t test (D) or Mann-Whitney test (C, G-K). Arrowheads indicate LGG puncta in the neurite. AIY boundary is indicated by dashed white line. Anterior is to the left.

### LGG-1 requires LGG-2 for puncta formation, but LGG-2 can compensate for the loss of LGG-1 in AIY

Previous studies suggest that LGG-1 and LGG-2 have both non-redundant and redundant functions, depending on the context (e.g. developmental stage, longevity, tissue type) (Alberti et al. 2010; Manil-Ségalen et al. 2014; Wu et al. 2015). However, many of these studies did not examine endogenous LGG-1 or LGG-2. Thus, we sought to examine whether LGG-1 or LGG-2 could compensate for the loss of the other at the pan-neuronal level. We first examined endogenous wrmNG_split_:LGG-1 puncta in *lgg-2(lof)* mutants. There were fewer, smaller, and dimmer wrmNG_split_:LGG-1 puncta in *lgg-2(lof)* mutants compared to wild-type animals (Fig. 5A-C). The significant decrease in the number of wrmNG_split_:LGG-1 autophagosomes suggests that LGG-2 is required for proper autophagosome formation. Further, these data indicate that LGG-1 does not compensate for a loss of *lgg-2* in neurons.

**Figure 5.**
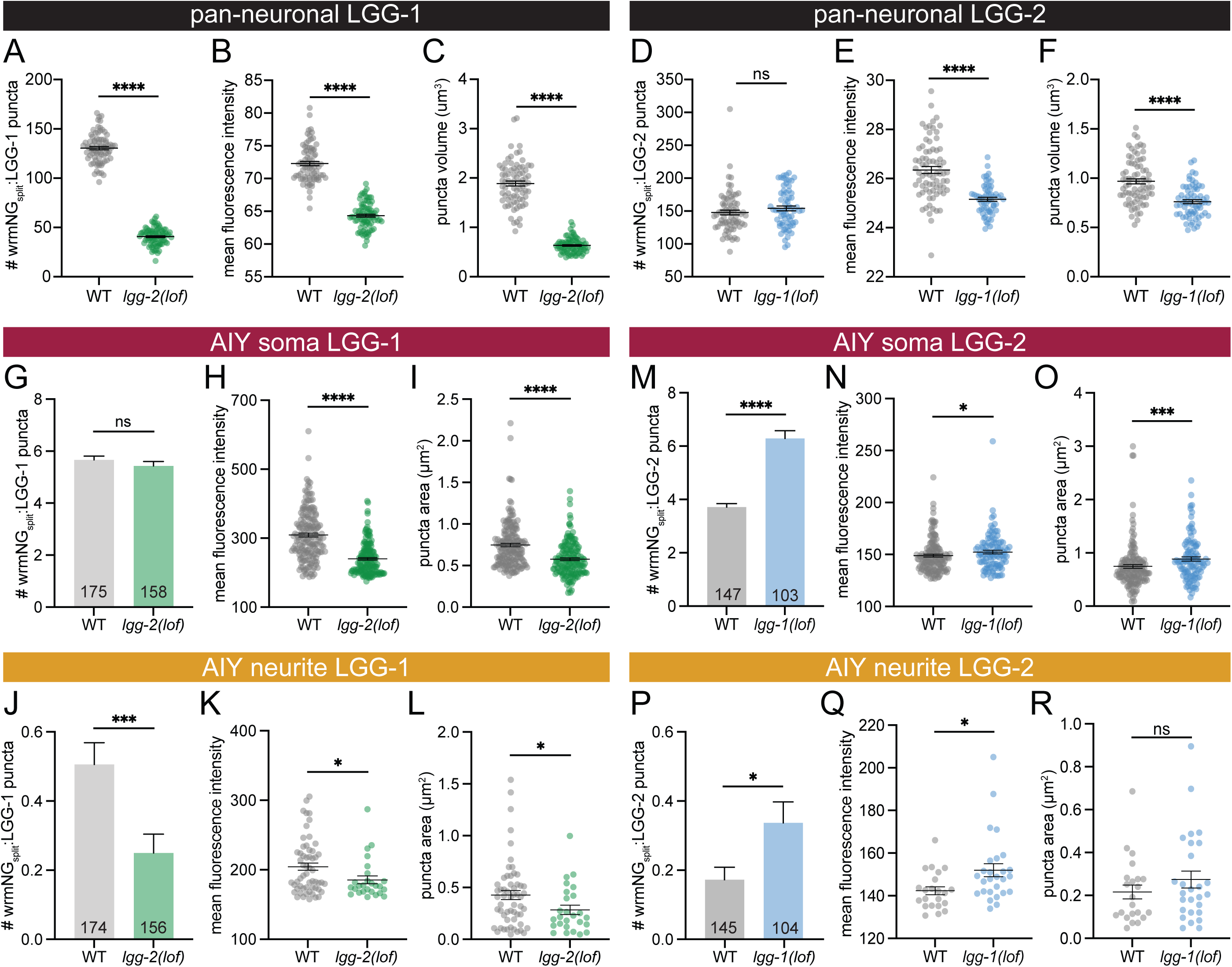
*lgg-2* is required for LGG-1 puncta formation and can compensate for the loss of *lgg-1* in AIY. (A-F) Quantification of wrmNG_split_:LGG-1 puncta in wild type and *lgg-2(lof)* mutant worms (A-C) and wrmNG:LGG-2 puncta in wild type and *lgg-1(lof)* mutant worms (D-F) in the nerve ring. Mean ± SEM. (G-L) Quantification of wrmNG_split_:LGG-1 puncta in wild type and *lgg-2(lof)* mutant worms in the AIY soma (G-I) and the AIY neurite (J-L). Mean + or ± SEM. (M-R) Quantification of wrmNG_split_:LGG-2 puncta in wild type and *lgg-1(lof)* mutant worms in the AIY soma (M-O) and the AIY neurite (P-R). Mean + or ± SEM. In G, J, M and P, n values are indicated at the bottom of each bar in the graph; in other graphs, all values are plotted. **** p<0.0001; *** p < 0.0005; * p < 0.05; ns p> 0.05 by Welch’s t test (A-B, E-G) or Mann-Whitney test (C-D, H-R).

Next, we quantified wrmNG_split_:LGG-2 puncta in *lgg-1(lof)* mutants to assess whether LGG-2 could compensate for the loss of *lgg-1*. We observed no difference in the number of wrmNG_split_:LGG-2 puncta between *lgg-1(lof)* mutants and wild-type worms (Fig. 5D), indicating that LGG-1 is not required for the formation of wrmNG_split_:LGG-2 puncta in neurons. However, *lgg-1(lof)* mutants also exhibited significantly dimmer (Fig. 5E) and smaller (Fig. 5F) wrmNG_split_:LGG-2 puncta compared to wild-type animals, suggesting that LGG-1 may be required for proper phagophore membrane expansion in neurons. Overall, these data indicate that LGG-1 and LGG-2 are both necessary for autophagosome formation, with no evidence of compensation upon the loss of either protein at the pan-neuronal level.

Because pan-neuronal analysis may mask changes in individual neurons, we next asked whether LGG-1 or LGG-2 could compensate for the loss of the other in AIY. First, we examined endogenous wrmNG_split_:LGG-1 puncta in *lgg-2(lof)* mutant animals (Fig. 5G-L). In these mutants, wrmNG_split_:LGG-1 puncta were smaller and dimmer than in wild-type animals in both the AIY soma (Fig. 5H-I) and the AIY neurite (Fig. 5K-L). The number of wrmNG_split_:LGG-1 puncta was not significantly different in the AIY soma (Fig. 5G), but the number of wrmNG_split_:LGG-1 puncta was reduced by 50% in the neurites of *lgg-2(lof)* mutant animals (Fig. 5J). These data reflect the pattern we observed at the pan-neuronal level and further reinforce that LGG-2 is required for LGG-1 puncta formation.

Next, we examined wrmNG_split_:LGG-2 puncta in AIY in *lgg-1(lof)* mutant animals (Fig. 5M-R). Interestingly, we detected a 69% increase in wrmNG_split_:LGG-2 puncta in the soma (Fig. 5M) and a 95% increase in wrmNG_split_:LGG-2 puncta in the neurite (Fig. 5P). These puncta were slightly, but significantly, brighter and larger in *lgg-1(lof)* mutants than in wild-type animals in both the AIY soma (Fig. 5N-O) and AIY neurite (Fig. 5Q-R). These data suggest that *lgg-2* is upregulated in AIY to compensate for the loss of *lgg-1.* This contrasts with what we observed at the pan-neuronal level, suggesting that certain neurons, such as AIY, may upregulate *lgg-2* to compensate for the loss of *lgg-1*, while others do not.

## DISCUSSION

Here we generated endogenously tagged *lgg-1* and *lgg-2*, the two *C. elegans* orthologs of Atg8, in the worm nervous system. We chose to engineer an endogenous tagging system that can be flexibly used to assess autophagy in individual tissues or even single neurons. Importantly, endogenous tagging of Atg8 in live animals removes overexpression artifacts while maintaining the ability to analyze the dynamics of the autophagy pathway. Further, a previous study used full-length fluorophores to endogenously tag *lgg-1* and *lgg-2* in *C. elegans* (Bördén et al. 2025). This strategy works well for tissues with large, easily identifiable cells such as the hypodermis or muscle. However, assessing autophagy with these reporters is incredibly challenging in the densely packed nerve ring with many neurites fasciculating together close to non-neuronal tissues. Furthermore, any differences in autophagy dynamics between neurons would be impossible to tease apart with existing endogenous autophagy reporters.

A previous study using single-cell spatial transcriptomics in *C. elegans* at the L4 stage demonstrated that *lgg-1* is more highly expressed than *lgg-2* in the nervous system (Taylor et al. 2021). Unsurprisingly, our endogenously labeled autophagy reporters mirrored these findings with wrmNG_split_:LGG-1 showing higher expression than wrmNG_split_:LGG-2 pan-neuronally and in individual AIY and NSM neurons. Overall, wrmNG_split_:LGG-1 puncta were also more numerous and larger than wrmNG_split_:LGG-2 puncta, consistent with their expression levels in neurons. Interestingly, a single neuron, I2, bucks the trend in the transcriptomics data, with *lgg-2* having slightly higher expression than *lgg-1* (Fig. 1B) (Taylor et al. 2021). It is intriguing to speculate that the trends we observed with endogenously labelled LGG-1 and LGG-2 in AIY and NSM would be reversed in I2.

Despite differences in expression levels between the worm Atg8s, we observed many similarities between endogenously labeled LGG-1 and LGG-2 puncta in the nervous system. Both populations of puncta were present in the same neuronal compartment, with autophagosomes labelled with either marker inhabiting the AIY soma and neurite and the NSM soma and axons. Both LGG-1 and LGG-2 puncta were also affected in autophagy mutants, with their endogenous signals dispersing into the cytoplasm. Additionally, UNC-16/JIP3 was required for retrograde transport of both wrmNG_split_:LGG-1 and wrmNG_split_:LGG-2 puncta (Fig. 4). Taken together, our data suggest that LGG-1 and LGG-2 might label overlapping populations or the same population of neuronal autophagosomes.

The roles of *lgg-1* and *lgg-2* in autophagy remain debated, with some reports suggesting distinct functions for each in autophagosome biogenesis and developmental bulk autophagy, while others support functional redundancy. We observed that, at a global nervous system scale, *lgg-1* and *lgg-2* play non-redundant roles in neuronal autophagy, as we did not observe a compensatory mechanism in *lgg-1(lof)* and *lgg-2(lof)* mutants. We found that *lgg-2* was required for wrmNG_split_:LGG-1 puncta formation in the nervous system broadly and specifically in AIY. We similarly determined that *lgg-1* was necessary for wrmNG_split_:LGG-2 puncta formation in NSM and pan-neuronally. However, when we analyzed wrmNG_split_:LGG-2 puncta specifically in AIY, we did not observe a similar decrease in LGG-2 puncta number, intensity, or area in *lgg-1(lof)* mutants. Together, these data suggest that while *lgg-2* is necessary for LGG-1 puncta formation, *lgg-1* is not required for the formation of LGG-2 puncta in AIY specifically. Despite this, we previously found that loss of either *lgg-1* or *lgg-2* phenocopies other autophagy mutants in AIY presynaptic assembly, indicating that both are required in AIY for proper autophagosome biogenesis (Stavoe et al. 2016).

Are wrmNG_split_:LGG-1 and wrmNG_split_:LGG-2 labelling the same population of puncta? Are there discrete populations of autophagosomes labelled by each Atg8 family? An obvious limitation of our system and this study is that we did not assess endogenously labeled LGG-1 and LGG-2 simultaneously. To do so, we needed two spectrally distinct split-fluorophore systems. A split-wrmScarlet system has been previously published (Goudeau et al. 2021). We continue to explore this and other systems to simultaneously visualize endogenous LGG-1 and LGG-2 in individual neurons.

While global analyses of autophagy across the nervous system are informative, our results underscore the necessity of systematic investigation of autophagy with single-neuron resolution. Our data show that pan-neuronal analyses can mask critical cell-specific differences. For example, in AIY, but not pan-neuronally, we observed an increase in number, intensity, and area of wrmNG_split_:LGG-2 autophagosomes in *lgg-1(lof)* mutants, suggesting that there is a capacity for redundancy between *lgg-1* and *lgg-2* in specific neurons. To our knowledge, this is the first report demonstrating this type of compensation. Further studies are necessary to determine how this compensatory mechanism functions in AIY and whether it extends to other individual neurons.

## ACKNOWLEDGEMENTS

The authors acknowledge the technical assistance of past Stavoe lab members Beverly Hughes and Ayesha Qureshi and helpful discussions and feedback from Arey lab members.

## METHODS

### C. elegans strains and maintenance

*C. elegans* strains were maintained at room temperature on NGM plates seeded with OP50 *E. coli(Brenner 1974)*. Some strains were provided by the *Caenorhabditis* Genetics Center (CGC), which is funded by NIH Office of Research Infrastructure Programs (P40 OD010440), including: N2 Bristol, WBM1215 *wbmIs89 [rab-3p::3xFLAG::dpy-10::SL2::wrmScarlet::rab-3 3’UTR, *wbmIs68]*, DRC4750 *olaIs35,* and CZ3011 *unc-16(ju146)*.

### Transgenic line generation

#### Split wrmNeonGreen

Optimized split mNeonGreen contains several point mutations compared to full-length mNeonGreen (Zhou et al. 2020). Codon-optimized wrmNeonGreen contains four synthetic introns (Hostettler et al. 2017). We used a combination of gene synthesis (IDT) and PCR-directed point mutagenesis to generate wrmNG3K_1-10_ (see supplemental data for sequence). wrmNG_11_ was generated by gene synthesis (IDT). We tested the split wrmNG system in AIY to ensure that the separate wrmNG_split_ fragments could complement in worm neurons to generate fluorescent signal before proceeding with endogenous tagging.

#### CRISPR-generated transgenic lines

An adapted co-CRISPR protocol (Ghanta and Mello 2020) was used to generate VOE211 *lgg-1(zny5[wrmNG11:lgg-1])*, VOE350 *lgg-1(zny5[wrmNG11:lgg-1]zny14[G116A])*, VOE514 *lgg-2(zny29[wrmNG11:lgg-2]),* VOE535 *lgg-1(zny35[A4*])*, VOE568 *lgg-2(zny29[wrmNG11:lgg-2]zny38[G130A])*, and VOE576 *lgg-2(zny37[P27*])*. For the co-CRISPR marker, the *dpy-10(cn64)* lesion, which produces worms with the roller or dumpy phenotype, or the sqt-2(sc3) lesion, which produces worms with the roller or squat phenotype, was recapitulated. All CRISPR reagents were designed with and obtained from Integrated DNA Technologies (IDT). CRISPR was performed in N2 worms, and edits were confirmed through Sanger sequencing. Successful lines were then outcrossed five times to N2 animals and confirmed to have no off-target edits by the CRISPR guide RNA and co-CRISPR guide RNA (as predicted by IDT) through Sanger sequencing before use in this study.

VOE287 *znyIs7(prab-3:wrmNG1-10:SL2:wrmScar:rab-3 3’UTR *wbmIs89)* animals were created using the SKI LODGE method (Silva-García et al. 2019). SKI LODGE CRISPR was performed following the same protocol as before but in WBM1215 worms using PCR-amplified wrmNG_1-10_ with homology arms as the repair template. Once again, edits were confirmed through Sanger sequencing. Successful lines were outcrossed five times to N2 animals and then confirmed to have no off-target edits.

#### Extrachromosomal arrays

Expression clones were created in the pSM vector (Shen and Bargmann 2003), a derivative of PD49.26 with extra cloning sites. Plasmid and transgenic strains were generated utilizing standard microinjection techniques and co-injected with punc-122:myr-mCh. The following plasmids were injected at the following concentrations: ptph-1:myr-mCh:SL2:wrmNG3K1-10:rab-3 3’UTR (20 ng/µL), pttx-3:wrmNG1-10:rab-3 3’UTR (30 ng/µL), pttx-3:egfp:lgg-2 (30 ng/µL), and pttx-3:mCh (30 ng/µL).

### Imaging and quantification

Prior to imaging, worms were mounted onto 2% agarose pads and immobilized in 10 mM levamisole (MilliporeSigma). Confocal Z-stack (step size: 0.3 um) images were captured using a spinning-disk confocal (Nikon Ti2 Inverted Confocal with Yokogawa W1 Spinning Disk Package) with Apochromat 60x or 100x 1.4 NA oil immersion objectives (Nikon Instruments). Digital micrographs were acquired with a back-illuminated cCMOS camera (Teledyne Photometrics) using Nikon Elements software. Multiple channels were acquired consecutively, with the green (488 nm) channel captured first, followed by the red (561 nm) channel. Imaging conditions (magnification, exposure, and laser power) were kept consistent across comparable strains.

All image analysis was performed on raw data. Nikon Elements was used to quantify LGG puncta in AIY on maximal intensity projections. LGG puncta in NSM were quantified using the FIJI analyze particles function on maximal intensity projections. Pan-neuronal LGG puncta were quantified using FIJI 3D objects counter. The FIJI Interactive 3D surface plot plugin was used to generate the surface plots depicted in Fig. 1D-I.

### Statistical analysis

Microsoft Excel was used for data assembly. Prism 10 (GraphPad) was utilized to perform statistical tests and to generate graphs. Data were tested for normality using the D’Agostino and Pearson test. Based on the normality analysis, the appropriate statistical test was selected.

Figure legends state the specific statistical tests utilized.

## KEY RESOURCES TABLE

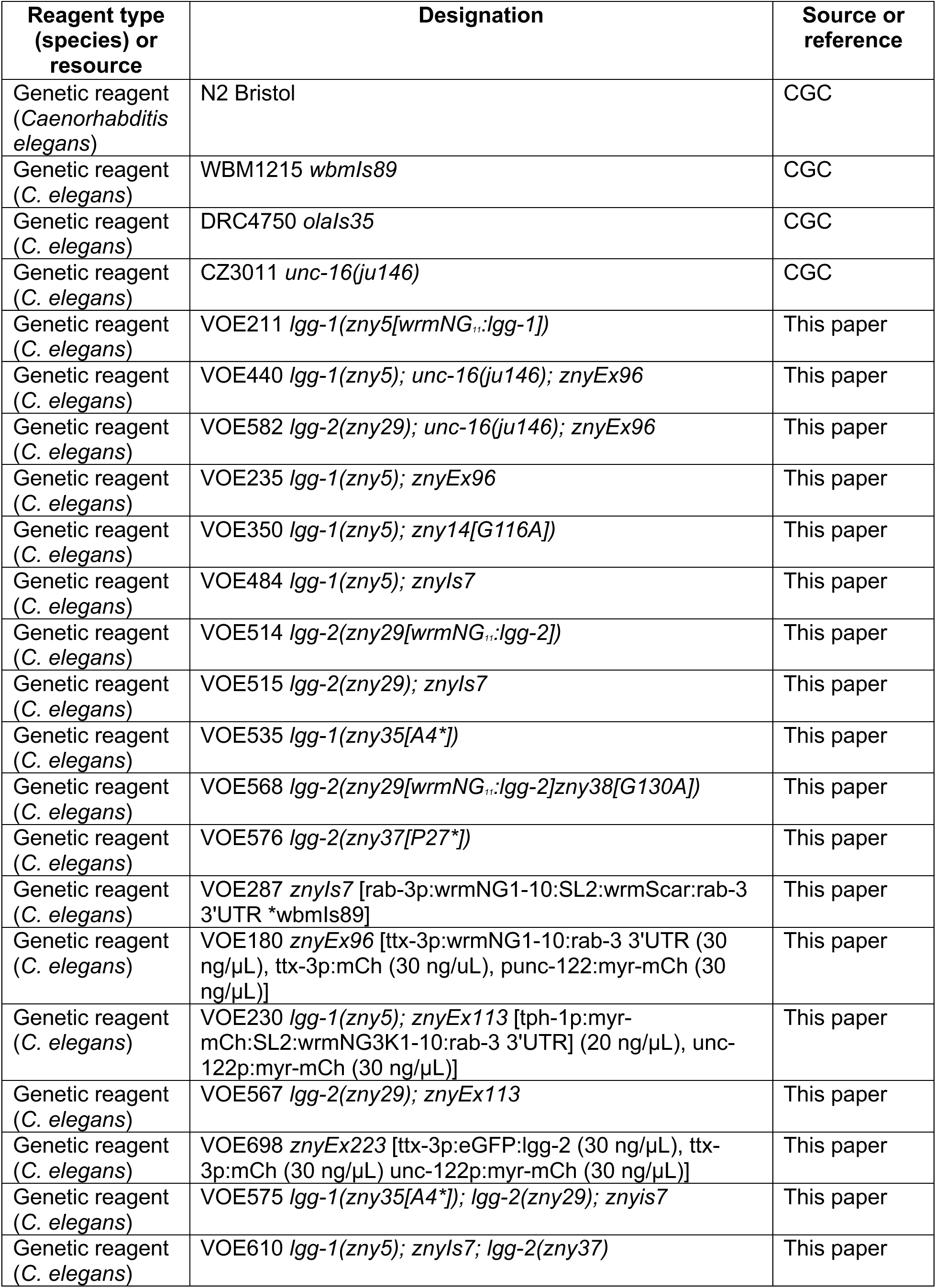
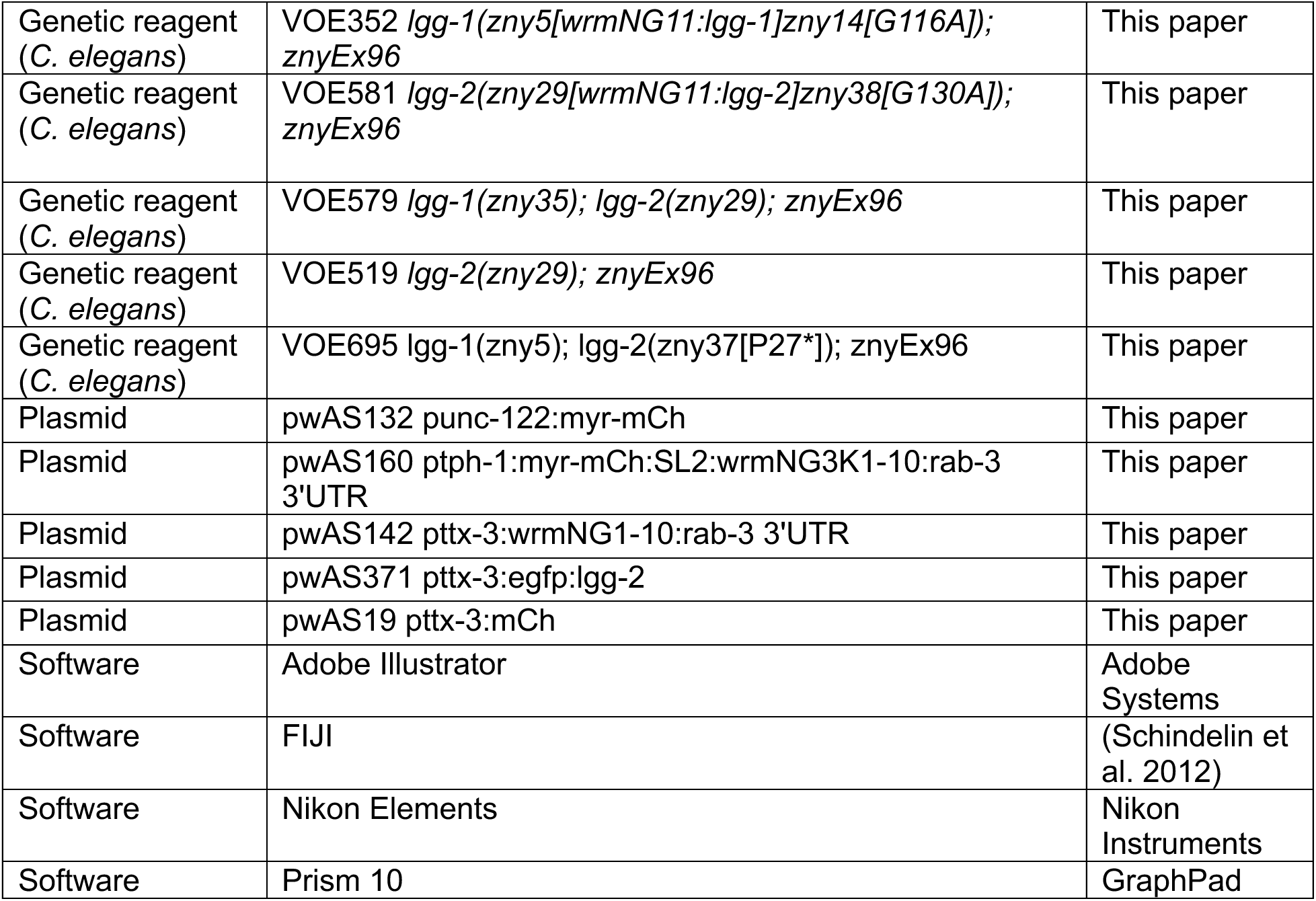

## SUPPLEMENTAL FIGURE LEGEND

**Figure S1.**
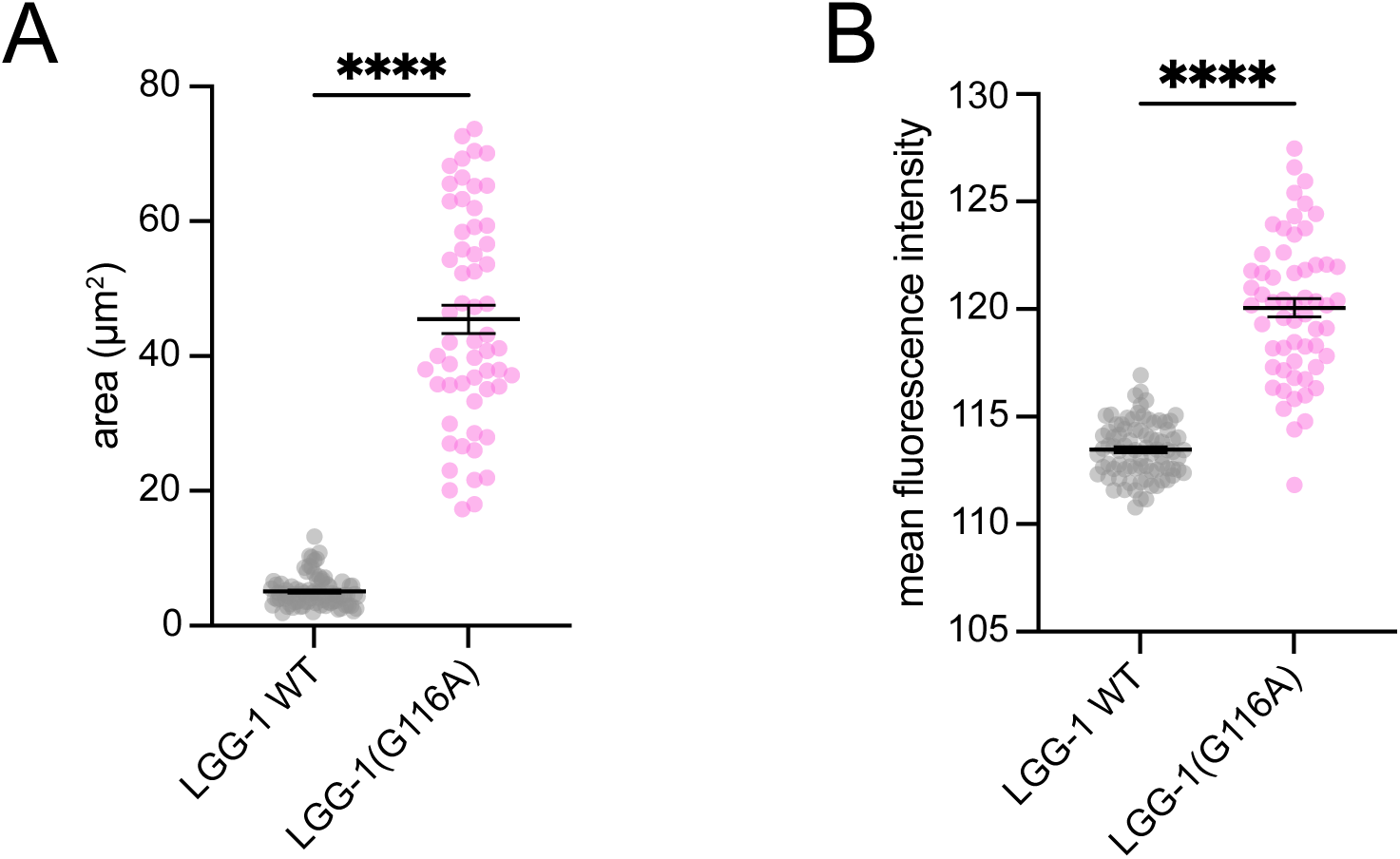
LGG-1(G116A) is brighter and more diffuse than wildtype LGG-1 in AIY. (A-B) Quantification of the area (A) and mean fluorescence intensity (B) of wrmNG_split_:LGG-1 and wrmNG_split_:LGG-1(G116A) signal, mean ± SEM, **** p<0.0001 by Welch’s t test.

## SUPPLEMENTAL DATA

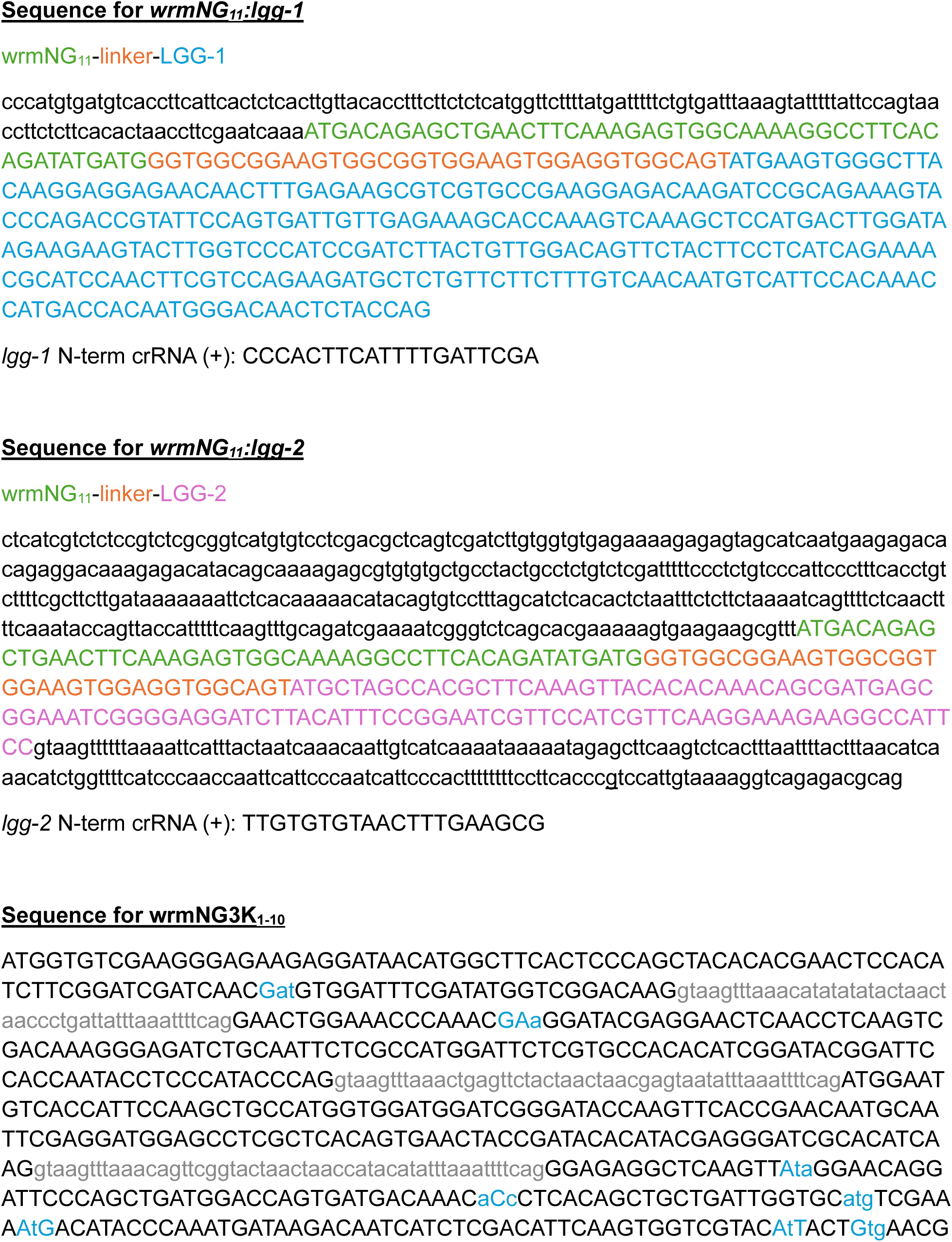
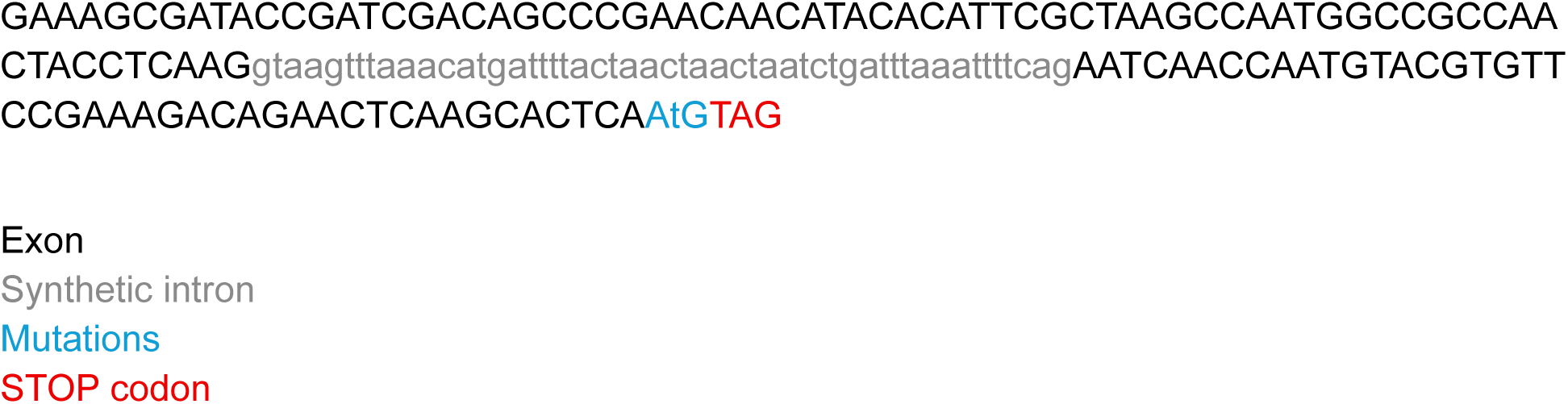

